# A genomic island of Streptomyces coelicolor with the self-contained regulon of an ECF sigma factor

**DOI:** 10.1101/247056

**Authors:** Camilla M. Kao, Nitsara Karoonuthaisiri, David Weaver, Jonathan A. Vroom, Shuning A. Gai, Mai-Lan Ho, Kedar G. Patel

## Abstract

Streptomycetes constitute the largest genus of actinobacteria, living predominantly in soil and decaying vegetation. The bacteria are widely known for their filamentous morphologies and their capacity to synthesize antibiotics and other biologically active molecules. More than a decade ago, we and others identified 22 genomic islands that *Streptomyces coelicolor* M145 possesses and other *Streptomyces* strains lack. One of these genomic islands, Genomic Island (GI) 6, encodes an extracytoplasmic function (ECF) sigma factor that we were characterizing in separate work. Here we report that artificial induction of the ECF sigma factor, which is encoded by SCO3450, causes the transcription of approximately one-fourth of GI 6, or ~26 mostly contiguous genes, to increase. More than half of the regulon encodes putative enzymes involved in small molecule metabolism. A putative haloacid dehalogenase is present. Genes encoding two putative anti-sigma factors flank SCO3450, the three genes residing within the regulon. Our data suggest that the ECF sigma factor and its regulon are a self-contained transcriptional unit that can be transferred by horizontal gene transfer. To our knowledge, only one other example has been identified of an ECF sigma factor and its contiguous regulon appearing to be transferrable by horizontal gene transfer [18,19]. Because the regulon appears not to be induced by the 44 growth conditions recently examined by Byung-Kwan Cho and colleagues [20], if it confers fitness to *S. coelicolor*, the regulon likely does so in as-yet unknown situations. Those situations might range from scavenging to detoxification to even communication within microbial communities.

**IMPORTANCE:** *Streptomyces* bacteria grow as hyphae that colonize soil and differentiate into spores when nutrients become scarce. In their terrestrial habitats, the bacteria encounter diverse conditions. Presumably so that the bacteria can cope with those conditions, the chromosomes of streptomycetes are highly dynamic, varying greatly in structure not only between species but also between closely related strains of a single species. The bacteria also have large numbers of extracytoplasmic function (ECF) sigma factors, which undoubtedly help the microorganisms respond to the plethora of challenges coming from the environment. This work illustrates these two threads of *Streptomyces* biology dovetailing: Genetic adaptability through horizontal gene transfer seems to have enabled *Streptomyces coelicolor* to acquire a self-contained transcriptional unit that consists of an ECF sigma factor and its regulon. The suggested facile movement of the regulon between microbial hosts indicates the value of the metabolism of small molecules possibly mediated by the regulon.

## Introduction

Researchers began studying *Streptomyces* genetics in the second half of the 20th century because of the filamentous morphologies and biosynthetic capacities of the bacteria [1]. Many of the *Streptomyces* strains that were studied belong to a group of closely related “blue” streptomycetes that had been given various names, but probably should all have the same name [2]. During the early 2000’s, we characterized the genomes of six of these “blue” strains by using DNA microarrays: the sequenced strain [3], *S. coelicolor* M145, which derives from *S. coelicolor* A3(2), which in turn derives from Waksman’s strain 3443 [4]; Sermonti’s SE1 (John Innes (JI) strain 1152) [5]; Bradley’s S199 (JI 1153) [6]; *S. lividans* 66 (JI 1326) [7]; *S. lividans* ISP5434 (JI 2896), which contains the plasmid pIJ101 [8]; and *S. violaceoruber* SANK95570 (JI 3034), which contains the plasmid pSV1 that encodes the biosynthetic genes for the antibiotic methylenomycin [9]. That study was described in the doctoral dissertation of co-author David Weaver [10]. Similar findings were reported by Jayapal et al., who examined *S. coelicolor* M145 and *S. lividans* TK21 [11].

By comparing the genome of *S. coelicolor* M145 to the five wild type genomes by using DNA microarrays, we identified 22 sets of contiguous genes in *S. coelicolor* M145 that are absent in the wild type strains [10]. We designated these genes Genomic Islands (GIs) 1 to 22. Fig. S1, Fig. S1, and Fig. S3 show the GIs. The sizes of the GIs range from 3 kb to 150 kb and the number of ORFs within them from 3 to 148. *S. coelicolor* M145 likely acquired the 22 GIs by horizontal gene transfer, because they have the following characteristics: Direct repeats flank most of the GIs; the GIs contain regions with low G + C content; the GIs encode transposable elements such as invertases, recombinases, and transposons, as well as plasmid-related proteins; some GIs appear to have inserted into tRNA sequences; and the GIs have slightly lower gene expression during exponential growth [10,11]. We found that different wild type strains lack different GIs and have different boundaries at the sites corresponding to the GIs in *S. coelicolor* M145. We also identified DNA that the wild type strains possess and that *S. coelicolor* M145 lacks. Some of that DNA (corresponding to GIs 8, 14, 17, 18, and 19 in *S. coelicolor* M145) might be large segments, because we were unable to amplify the DNA by PCR [10]. The genomic islands have a relatively uniform distribution across the chromosome of *S. coelicolor* M145, with a slight abundance in the right chromosome arm [10,11].

A sigma factor is a specialized unit of RNA polymerase that permits the multisubunit enzyme to initiate transcription selectively. Extracytoplasmic function (ECF) sigma factors constitute the most abundant, smallest, and most divergent group of sigma factors [12]. Their name derives from many of their members having roles in sensing and responding to signals generated outside of the cell or in the cell membrane [13]. The genome of *S. coelicolor* M145 encodes 63 sigma factors, of which 49 belong to the ECF group [3]. During the early 2000’s, we sought to identify regulons of *S. coelicolor* sigma factors by overexpressing each one individually and analyzing the response of the transcriptome by using DNA microarrays. The ECF sigma factor encoded by SCO3450 was evaluated as part of that work. While we were unaware of this fact at that time, SCO3450 is located in GI 6 [10].

The ECF sigma factor encoded by SCO3450 belongs to the subgroup ECF01 as classified by Thorsten Mascher and colleagues in 2009 [14]. This subgroup includes RpoE-like sigma factors, of which the best studied is σ^W^ of *Bacillus subtilis*. Members of the ECF01 subgroup are widely distributed throughout bacterial phyla, but are absent in most actinobacteria, the phylum which includes the genus *Streptomyces* [15]. ECF01 sigma factors have been characterized experimentally to be involved in responses to envelope stress and the production and detoxification of antimicrobial compounds [14].

## Results and Discussion

### Genomic Island 6 encodes an ECF sigma factor and the regulon of the sigma factor

Table 1 lists the five wild type *Streptomyces* strains that we studied that lack GI 6 [10].

**Table 1.** Strains that revealed the genomic islands of *S. coelicolor* M145.

| Strain | Comment | Ref. |
| --- | --- | --- |
| M145 | *S. coelicolor* sequenced "reference" strain, SCP1- SCP2- | [3] |
| 1152 | *S. coelicolor* Sermonti's SE1 | [5] |
| 1153 | *S. coelicolor* Bradley's S199 strain | [6] |
| 1326 | *S. lividans* 66 | [7] |
| 2896 | *S. lividans* ISP5434-, the strain from which pIJ101 was isolated | [8] |
| 3034 | *S. violaceoruber* SANK95570, harbors pSV1, which encodes methylenomycin biosynthetic genes | [9] |

Fig. 1 shows the boundaries of GI 6 (the left-hand columns) and the levels of transcripts, as measured by microarrays, after artificial induction of SCO3450 in the strain *S. coelicolor* M600 growing exponentially in a liquid culture (the right-hand columns) [16]. Fifteen minutes after we induced the sigma factor, transcripts of approximately one-fourth of the genes in GI 6 increased in abundance. The genes numbered ~26: SCO3437 and SCO3442-3465; SCO3478 is an induced gene located 14 kb away from the other genes. Cross-hybridization does not explain the result for SCO3478 because the gene, which encodes a dehydrogenase, lacks sequence similarity to other *S. coelicolor* genes. SCO3450 is located among the induced genes (Fig. 1, arrow; Fig. S4). Together these data suggest that the ECF sigma factor and its regulon are a self-contained transcriptional unit that can be transferred by horizontal gene transfer.

**FIG 1.**
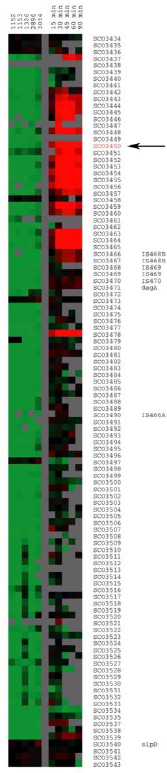
Genomic Island 6 and the regulon of the ECF sigma factor encoded by SCO3450. Genomic Island 6 and the regulon of the ECF sigma factor encoded by SCO3450. GI 6 is 108 kb in length. The arrow indicates the location of SCO3450, which encodes a putative ECF sigma factor. See Table 1 for a list of the strains used to identify GI 6. To obtain the data in the figure, we used DNA microarrays that contained the 7825 predicted genes in the chromosome of *S. coelicolor* M145 [17]. For the left-hand columns, the maximum intensity of green corresponds to a 3-fold difference in signal on the microarrays for DNA in *S. coelicolor* M145 relative to DNA in the wild type strains. SCO3450 was overexpressed in an exponentially growing liquid culture (SMM) of a *S. coelicolor* M600 derivative, through the use of a thiostrepton-inducible promoter [16]. RNA was harvested from the culture immediately prior to the addition of thiostrepton and every 15 to 30 minutes afterwards for 90 minutes. cDNA of the initial RNA sample (OD_450_ ~ 0.5) was labeled with the green fluorescent dye Cy3. cDNA of RNA isolated from subsequent time points was labeled with the red fluorescent dye Cy5. Yellow microarray spots for a particular time point represented genes with transcript levels equal to the levels of the initial time point. Red and green microarray spots represented genes induced and repressed, respectively, after the addition of thiostrepton. The yellow color is shown here as black for clarity. For the right-hand columns, the maximum intensity of red corresponds to 8fold induction of a given gene at a particular time relative to the beginning of the time course. A control experiment with a M600 derivative that lacks SCO3450 in the induction plasmid identified genes induced by thiostrepton. Labels on the right indicate ORF numbers and genes with names.

To our knowledge, the only other example of an ECF sigma factor and its regulon appearing to be transferrable by horizontal gene transfer is the system for cobalt and nickel resistance in the bacterium *Cupriavidus metallidurans* CH34 [18,19]. There, the plasmid pMOL28 contains six *cnr* genes that are organized in two adjacent operons of three genes each. The gene *cnrH* encodes the ECF sigma factor CnrH. Deletion and complementation of *cnrH* indicated that CnrH is essential for the regulation of the *cnr* system. When CnrH was present but the periplasmic sensor CnrX and the anti-sigma factor CnrY were absent, high-level constitutive expression was observed for the *cnr* genes [18].

### The regulon of the ECF sigma factor contains conserved putative −35 and −10 promoter regions

Byung-Kwan Cho and colleagues reported in 2016 transcription start sites of *S. coelicolor* genes induced by 44 growth conditions [20]. Those conditions included growth in rich media and many kinds of minimal media, growth in the liquid phase and on solid media, rapid growth to stationary phase, and several kinds of shocks. That study did not identify transcription start sites for the regulon of the ECF sigma factor encoded by SCO3450. This observation indicates that the regulon is not induced by the growth conditions examined by Jeong et al. [20]. Perhaps *S. coelicolor* uses the regulon for circumstances that are not replicated in the laboratory, such as interactions in the natural environment among different microbial species.

We used the bioinformatics tool PromoterHunter [21], combined with visual inspection of nucleotide sequences, to identify possible −35 and −10 promoter regions in the regulon of the ECF sigma factor. The conserved sequences for the −35 and −10 promoter regions might be AACGG and CG, respectively (Fig. 2). The promoters of 14 genes contain these sequences exactly. The data suggest that operons might comprise SCO3445 to SCO3444, SCO3448 to SCO3446, SCO3455 to SCO3452, and SCO3463 to SCO3465 (see Fig. S4). Alternatively, genes not listed in Fig. 2 might have promoters with less conserved −35 and −10 regions. Note that SCO3460 and SCO3478 do not possess the highly conserved promoter regions (see Fig. S4). In addition, that SCO3438 has the conserved promoter regions supports the notion that this gene belongs to the regulon of the ECF sigma factor. In Fig. 1, the lack of data for SCO3438 from all of the microarray hybridizations indicates that its spot on the microarrays was functioning poorly.

**FIG 2.**
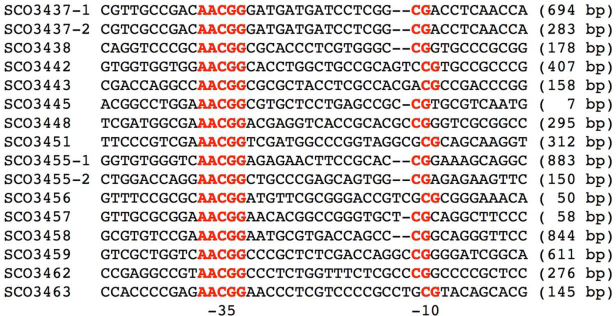
Alignment of putative promoters in the regulon of the ECF sigma factor encoded by SCO3450. Alignment of putative promoters in the regulon of the ECF sigma factor encoded by SCO3450. Red letters indicate the putative −35 and −10 promoter regions. On the right-hand side, the numbers in parentheses indicate the distance between each −10 promoter region and the start codon of the respective gene.

### The regulon of the ECF sigma factor encodes putative enzymes for small molecule metabolism

Recently we updated the annotations of the regulon of the ECF sigma factor by using EnsemblBacteria [22] and UniProt BLASTS [23]. Of the approximately 26 genes in the regulon, 15 encode putative enzymes for small molecule metabolism. 11 genes encode putative membrane and transport proteins. Four genes encode putative regulatory proteins. Four of the 26 genes fall into two categories. Only one gene lacks a putative function. Table S1 lists the annotations of each gene in the regulon.

### Enzymes encoded by the regulon of the ECF sigma factor

Table 2 lists the types of enzymes encoded by the regulon of the ECF sigma factor. The table excludes SCO3461 and SCO3462, because they lack gene expression data (Fig. 1, right-hand columns).

**Table 2.** Types of enzymes encoded by the regulon of the ECF sigma factor.

| Type | Gene | AA | Putative Function | Other feature |
| --- | --- | --- | --- | --- |
| **CLEAVAGE** |  |  |  |  |
| Glycoside hydrolase | SCO3444 | 617 | glycoside hydrolase family 15/phosphorylase b kinase regulatory chain family; six-hairpin glycosidase |  |
| Nucleoside phosphorylase | SCO3463 | 262 | nucleoside phosphorylase domain |  |
| Dehalogenase | SCO3446 | 225 | haloacid dehydrogenase (HAD)-like domain |  |
| **REDOX** |  |  |  |  |
| Glutaredoxin | SCO3442 | 114 | glutaredoxin (DNA synthesis?) |  |
| Oxidoreductase | SCO3443 | 454 | pyridine nucleotide-disulphide oxidoreductase; dihydrolipoyl dehydrogenase? |  |
|  | SCO3460 | 505 | pyridine nucleotide-disulphide oxidoreductase; dihydrolipoamide dehydrogenase? |  |
| Dehydrogenase | SCO3478 | 344 | D-isomer specific 2-hydroxyacid dehydrogenase |  |
| **TRANSFER** |  |  |  |  |
| Methyltransferase | SCO3452 | 359 | S-adenosyl-L-methionine-dependent methyltransferase |  |
|  | SCO3459 | 287 | S-adenosyl-L-methionine-dependent methyltransferase |  |
| Phosphatidyltransferase? | SCO3457 | 205 | CDP-alcohol phosphatidyltransferase; YnjF? | transmembrane |
| Nucleotide-diphospho-sugar transferase | SCO3464 | 210 | nucleotide-diphospho-sugar transferases; transferase 1, rSAM/selenodomain-associated |  |
|  | SCO3465 | 236 | nucleotide-diphospho-sugar transferase; transferase 2, rSAM/selenodomain-associated |  |
| **MISC** |  |  |  |  |
| Enzyme? | SCO3445 | 55 | low to moderate homology to small regions of seven larger proteins, which include enzymes | membrane |

AA = amino acids

The putative enzymes fall into three general groups. One group possibly catalyzes the cleavage of small molecules. The group includes a glycoside hydrolase, a nucleoside phosphorylase, and a haloacid dehalogenase. A second group of enzymes might catalyze oxidation and reduction reactions. The group includes a glutaredoxin-like protein, two oxidoreductases, and a dehydrogenase. The third group of enzymes might catalyze transfer reactions. The group includes two methyltransferases, perhaps a phosphatidyltransferase, and two nucleotide-diphospho sugar transferases. A final gene, SCO3445, encodes a small membrane protein that has 30-50% identity to small regions of seven larger proteins, a set which includes several enzymes.

The composition of the enzymes in the regulon of the ECF sigma factor suggests that the regulon might help to metabolize small molecules. A mixture of compounds might be being degraded: sugars, nucleosides and nucleotides, and lipids for the purposes of energy and biosynthesis; halogenated molecules for detoxification. Because the regulon encodes a set of putative enzymes with diverse substrates, the small molecules might be coming from the environment, for example, from neighboring cellular compartments or hyphae that are lysing due to hostile conditions.

Haloacid dehalogenases (HADs) belong to a large superfamily of hydrolases with diverse substrate specificity [24]. Type II HADs catalyze the hydrolytic dehalogenation of small L-2-haloalkanoic acids to yield the corresponding D-2-hydroxyalkanoic acids [25]. Because many *Streptomyces* bacteria produce halogenated antibiotics [26,27], the dehalogenase encoded by SCO3446 might serve to defend *S. coelicolor* against competitors in the environment by helping to catabolize antibiotics.

The following pairs of genes have no significant similarity, such that no cross-hybridization on the microarrays should have occurred: SCO3452 and SCO3449, which encode the methyltransferases; SCO3443 and SCO3460, which encode the oxidoreductases; and SCO3464 and SCO3465, which encode the nucleotide-diphospho-sugar transferases (Table 2).

### Anti-sigma factors within the regulon of the ECF sigma factor

Two genes that possibly encode anti-sigma factors flank SCO3450, the gene encoding the ECF sigma factor. The gene product of SCO3451 encodes a 103-amino-acid protein. A UniProt BLAST shows that the protein has approximately 40% identity with varying coverages to putative anti-sigma factors of other bacterial species, including putative transmembrane anti-sigma factors. The Constrained Consensus TOPology (CCTOP) prediction server [28] predicts with a reliability of 95.9069 that SCO3451 encodes a protein with one transmembrane segment (Fig. S5). The gene product of SCO3451 also contains a putative zinc-finger found in some anti-sigma factor proteins (Fig. S6). This zinc finger domain overlaps with the predicted transmembrane domain.

SCO3449 encodes a 106-amino-acid protein. A UniProt BLAST shows that the protein has approximately 40% identity with coverages around 50% to putative anti-sigma factors in other bacterial species, including putative transmembrane anti-sigma factors. However, according to CCTOP, the protein does not contain a transmembrane domain. Like the putative anti-sigma factor encoded by SCO3451, the putative anti-sigma factor encoded by SCO3449 contains a possible zinc-finger found in some anti-sigma factor proteins (Fig. S7).

Microarray data are lacking for the expression of SCO3449 (Fig. 1, right-hand columns), such that inferences about the role of this gene in comparison to those of the induced genes should be made with caution. However, because microarray data are available for this gene in the hybridizations that identified the genomic islands (Fig. 1, left-hand columns), the spot on the microarrays that represented this gene was likely intact. It is possible that a low abundance of transcripts of SCO3449 in both the reference and experimental samples of mRNA produced poor signals from the microarray spot of the gene.

### Membrane and transport proteins encoded by the regulon of the ECF sigma factor

Four contiguous genes in the regulon of the ECF sigma factor, SCO3453 to SCO3456, encode proteins that likely constitute an ABC transporter system. UniProt BLASTS show that all of the proteins have 45-50% identity with coverages greater than 95% to homologs with putative functions in spermidine and putrescine transport in other bacterial species. In particular, the species include *Geodermatophilus* and *Wenxinia*. Spermidine is a polyamine involved in cellular metabolism that can be used to stimulate RNA polymerase. Putrescine attacks S-adenosyl methionine and converts it to spermidine [29].

The putative transporters Sco3454 and Sco3455 have 34% identity with each other, the coverage being 44% between the C-terminal portions of the proteins.

### Genomic Island 6 consists of four segments with transposases at their boundaries

In addition to the regulon of the ECF sigma factor, we updated the annotations of the other genes of Genomic Island 6 by using EnsemblBacteria and UniProt BLASTS (Table S2). In the table, the location of the regulon is denoted by the words “REGULON HERE.”

The length of GI 6 is 108 kb. The genomic island consists of four segments of DNA bounded by transposases (Table S2, pink color). Three of the four segments have coherent putative functions: the oxidation and reduction of copper (Table S2, blue color); a characterized agarase encoded by *dagA* [30,31]; and the utilization of sugars (Table S2, yellow color). Included in the latter segment are a putative *lacI*-family transcriptional regulator and a putative β-galactosidase. A large fourth segment encodes putative enzymes and many hypothetical proteins.

The coherent functions of at least three of the four segments of GI 6 indicate that significant portions of the island might be “active.” If so, the regulon of the ECF sigma factor is likely active as well. While a nucleotide blast of the regulon of the ECF sigma factor yielded no similar segments of DNA in other sequenced organisms, the value to *S. coelicolor* of the regulon for as-yet unknown reasons is indicated by the presence of the regulon on a genomic island. It would be interesting to determine how common the regulon is among natural populations of microbes and whether the regulon moves easily between hosts.

## Materials and Methods

### *Construction of a* S. coelicolor *strain that overexpresses SCO3450*

SCO3450, the gene which encodes the ECF sigma factor harbored by GI 6, was amplified by PCR and cloned into the conjugative plasmid pIJ6902 [32] by using the restriction sites *NdeI* and

*Bgl*II. These restriction sites placed the gene immediately downstream of the thiostrepton-inducible promoter *tipAp*. The resulting plasmid was transformed into the methylation-deficient strain ET12567 with a non-transmissable helper plasmid, pUZ8002, and conjugated into *S. coelicolor* M600 as described by Kieser et al. [33]. The exconjugants were selected by an overlay of 50 µg/mL of apramycin.

### Time course of the overexpression strain

Supplemented minimal medium (SMM) was inoculated with approximately 5 × 10^7^ spores/mL of a spore stock. The culture was grown at 30°C to early exponential phase, which corresponded to a cell density of OD450 ~ 0.5. A reference sample, designated “0 min,” was harvested. To induce transcription of SCO3450, thiostrepton was added to the culture to a final concentration of 30 µg/mL. Samples of the culture were harvested 15, 30, 45, 60, and 90 minutes after induction of SCO3450.

### Extraction of total RNA from the overexpression strain

Samples of cells from liquid cultures were recovered by filtration on Whatman filter paper (15-20 mm diameter; catalog #1002 055). RNA was isolated using the modified Kirby mix protocol as described previously [34] with the following modifications: Harvested volumes ranged between 5 mL and 20 mL, depending on the cell densities. RNA samples were treated only once with DNase I (50-70 units; RNase-free, Invitrogen) for 15 minutes at room temperature.

### DNA microarray experiments for the overexpression strain

Samples of cDNA were synthesized from total RNA as described previously [34]. cDNA of the reference sample was labeled with Cy3-CTP. cDNAs of the samples isolated at subsequent time points were labeled with Cy5-CTP. The cDNAs were hybridized to microarrays as described previously [34].

### Identification of −35 and −10 conserved promoter regions in the regulon of the ECF sigma factor

The tool PromoterHunter [21] was used to examine DNA upstream of the genes in the regulon of the ECF sigma factor. Weight matrices corresponding to AAC for the −35 promoter region and CG for the −10 promoter region were used, because they reflect the conserved promoter regions of the subgroup ECF01 of ECF sigma factors [14]. A global G + C content of 72% was used. The space between the −35 and −10 regions was specified to be between 17 to 21 base pairs. Initially DNA segments of 300 base pairs were examined for isolated genes and genes located at the beginning of likely operons. Visual inspection of sequences returned by PromoterHunter identified AACGG and CG as possible consensus sequences for the −35 and −10 promoter regions, respectively. PromoterHunter was used to search for these conserved sequences within DNA segments of 1000 base pairs upstream of all of the genes in the regulon, in order to ensure that all instances of the sequences were identified.

## Acknowledgements

We thank Thomas M. Privalsky for informing us about bioinformatic tools and Mark J. Buttner and David A. Hopwood for discussions about the data.

## Author Contributions

C.M.K. conceived the experiments. D.W. designed the experiments that compared the genomes of the six *Streptomyces* strains and performed those experiments with J.A.V., M.L.H., and K.G.P. N.K. designed the experiment overexpressing the ECF sigma factor and performed the experiment with S.A.G. C.M.K. conducted the recent bioinformatic analyses and wrote the manuscript.

## Legends for Supplemental Material

**FIG S1** Genomic Islands 1 to 10 of *S. coelicolor* M145. Only the genomic islands are shown and not the entire chromosome. Table 1 lists the strains used. To obtain these data, we used DNA microarrays that contained the 7825 predicted genes in the chromosome of *S. coelicolor* M145 [17]. Genomic DNA from *S. coelicolor* M145 was used as the reference sample and labeled with the green fluorescent dye Cy3. Genomic DNA samples from the five wild type strains each were labeled with the red fluorescent dye Cy5. The labeled DNA of each wild type strain was mixed with the labeled DNA of *S. coelicolor* M145 and hybridized to a microarray. Yellow spots on the microarrays represented genes at equal copy numbers between *S. coelicolor* M145 and the other strains. Green spots represented genes present in *S. coelicolor* M145 but absent in the other strains. Red spots represented genes at higher copy numbers in the strains relative to *S. coelicolor* M145. The yellow color is shown here as black for clarity. The maximum intensity of green corresponds to a 3-fold difference in signal on the microarrays for DNA in *S. coelicolor* M145 relative to DNA in the wild type strains. The green bars to the left of the GIs designate horizontally transferred genes in the genome sequence of *S. coelicolor* M145 that were predicted by Bentley et al. [3]. Regions of GIs not predicted by Bentley et al. lack a bar. Labels on the right indicate ORF numbers and genes with names.

**FIG S2** Genomic Islands 11 to 18 of *S. coelicolor* M145. See the text of Fig. S1.

**FIG S3** Genomic Islands 19 to 22 of *S. coelicolor* M145. See the text of Fig. S1.

**FIG S4** The region of the *S. coelicolor* M145 chromosome that contains the regulon of the ECF sigma factor. The figure was obtained from EnsemblBacteria [22]. SCO3478 is not shown.

**FIG S5** Result from CCTOP for the putative anti-sigma factor encoded by SCO3451. The putative anti-sigma factor is predicted to contain one transmembrane domain. The data were obtained from the Constrained Consensus TOPology (CCTOP) prediction server [28].

**FIG S6** Predicted zinc finger in the putative anti-sigma factor encoded by SCO3451. The data were obtained from UniProt [23].

**FIG S7** Predicted zinc finger in the putative anti-sigma factor encoded by SCO3449. The data were obtained from UniProt [23].

**Table S1. Annotations of the regulon of the ECF Sigma Factor encoded by SCO3450.** Annotations were obtained from EnsemblBacteria [22] and UniProt [23]. For the UniProt BLASTs, parentheses denote genera and percent identities of similar proteins. Red text denotes putative enzymes. Blue text denotes proteins with homology to small regions of larger proteins.

**Table S2. Annotations of Genomic Island 6, excluding the regulon of the ECF sigma factor encoded by SCO3450.** Annotations were obtained from EnsemblBacteria [22] and UniProt [23]. For the UniProt BLASTs, parentheses denote genera and percent identities of similar proteins. Red text denotes putative enzymes. Blue text denotes proteins with homology to small regions of larger proteins. See the text for additional details.

